# Aegeline and Atorvastatin Synergistically Attenuate oxLDL-Induced Inflammation and Intracellular Cholesterol Accumulation in THP-1 Macrophages

**DOI:** 10.64898/2026.08.14.744794

**Authors:** Abinayaa Rajkumar, Cathline Mary Ramesh, Ezhila Subramani Dhatchana moorthy Vedhanayaki, Kalaiselvi Periandavan

## Abstract

**Background:** Atherosclerosis is driven by macrophage foam cell formation resulting from excessive oxidized low-density lipoprotein (oxLDL) accumulation and chronic vascular inflammation. This study evaluated the therapeutic potential of Aegeline, Atorvastatin, and their combined in mitigating oxLDL-induced inflammatory responses, cholesterol accumulation, and oxLDL uptake in human THP-1 macrophages.

**Methods:** THP-1 monocytes were differentiated into macrophages using a 72-hour differentiation protocol followed by a 48-hour resting period, confirmed via CD14 surface marker characterization. Macrophages were exposed to DiI-oxLDL and treated with Aegeline, Atorvastatin, or their combination. Key inflammatory cytokines and chemokines (CRP, TNF-α, IL-6, and IL-8) were measured using ELISA. Cholesterol efflux capacity and cellular oxLDL uptake were quantitatively assessed using fluorescence retention assays and immunofluorescence imaging.

**Results:** Differentiation of THP-1 monocytes to macrophages resulted in marked down-regulation of CD14 expression. DiI-oxLDL exposure triggered significant pro-inflammatory mediator secretion (p<0.001) and excessive intracellular cholesterol accumulation. Single-agent treatment with Aegeline or Atorvastatin significantly attenuated oxLDL-induced elevations of CRP, TNF-α, IL-6, and IL-8. Atorvastatin alone strongly suppressed CRP expression back to physiological baseline levels (p=ns vs. control). Notably, the combination of Aegeline and Atorvastatin demonstrated enhanced, broad-spectrum anti-inflammatory efficacy, achieving superior suppression of TNF-α (p=ns vs. control), IL-6, and IL-8 compared to monotherapies. Furthermore, both agents promoted cholesterol efflux and suppressed oxLDL uptake, with the combination treatment producing the lowest residual intracellular cholesterol levels (p<0.001).

**Conclusion:** Aegeline and Atorvastatin effectively suppress oxLDL-induced macrophage inflammatory cascades and intracellular lipid overload. While Atorvastatin monotherapy exerts robust control over CRP and oxLDL loading, combining Aegeline with Atorvastatin provides synergistic efficacy, enhancing cholesterol efflux and restoring pro-inflammatory cytokine expression toward physiological levels.

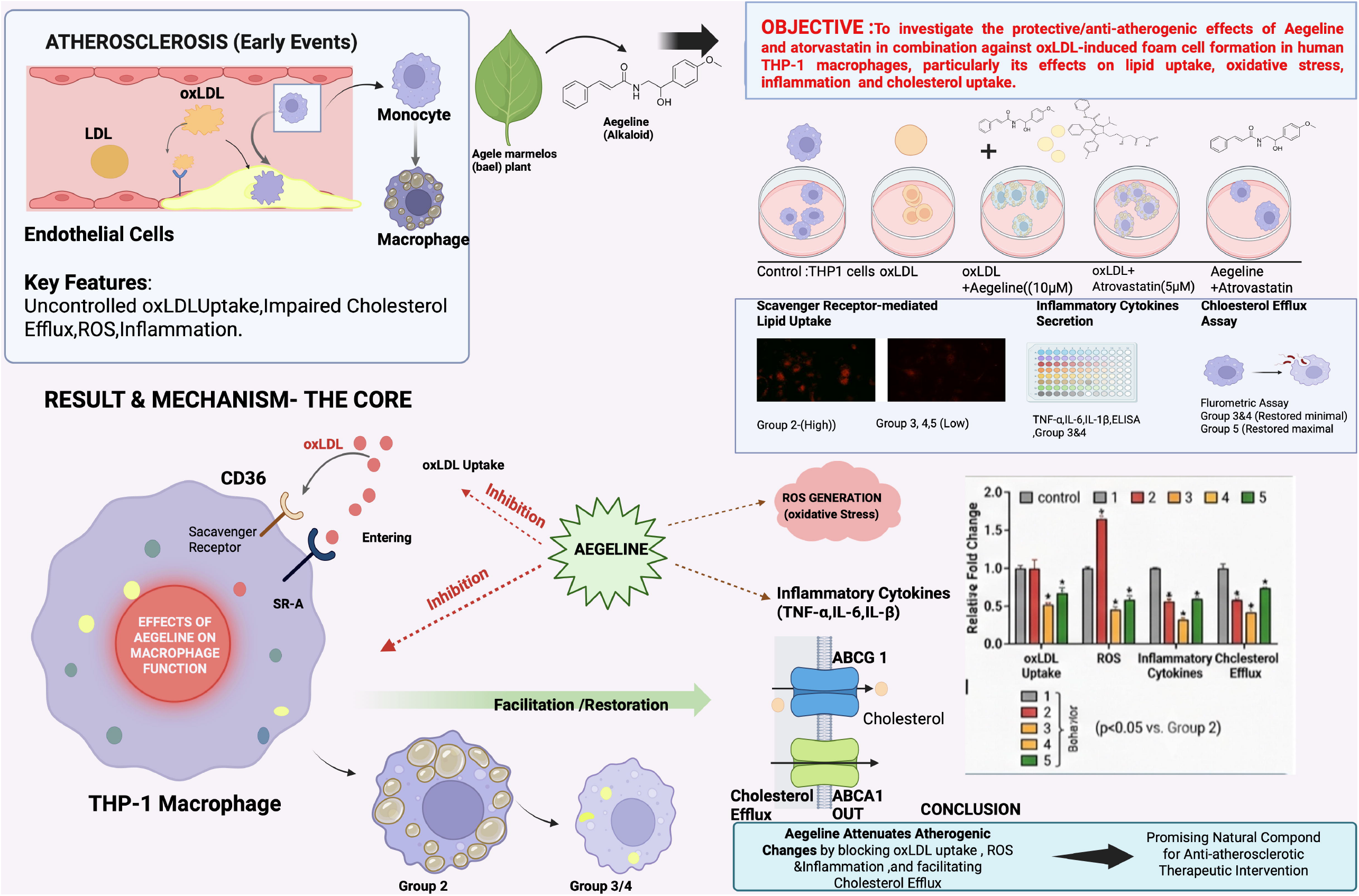

## 1.0 Introduction

Atherosclerosis remains the leading pathological substrate underlying cardiovascular disease and foremost cause of morbidity and mortality globally. This is initiated by dysfunction of endothelium and subendothelial retention of low-density lipoprotein (LDL), that subsequently undergoes oxidative modification of LDL to form oxidized LDL (oxLDL) (Borén *et al*., 2025) Unlike native LDL, oxLDL is avidly recognized and internalized by monocyte-derived macrophages through scavenging receptors such as SR-A, CD36 and LOX-1 (Steinberg D *et al*., 1995). The uptake of lipid eventually ends up with progressive accumulation of intracellular cholesteryl esters, macrophages reprogramming into lipid-laden “foam cells”, the defining traditional hallmark of the early atherosclerotic lesion (Leiviskä *et al*., 2013). Foam cell formation is a dynamic and self-amplifying pathological process. Intracellular oxLDL accumulation triggers a surge in reactive oxygen species (ROS) generation, which in turn promotes further oxidation of LDL, activates redox-sensitive transcription factors such as NF-κB, and amplifies oxidative stress. This oxidative burden is tightly coupled with pro-inflammatory macrophage phenotype characterized by enhanced cytokine secretion includes tumor necrosis factor-alpha (TNF-α) (Kleinbongard *et al*., 2010) interleukin-6 (IL-6), and interleukin-8 (IL-8), together, these events fuel inflammation, recruit additional monocytes, and destabilize the growing plaque. In parallel, cholesterol transporters facilitate the excess intracellular cholesterol to extracellular acceptors like apolipoprotein A-I and HDL, become functionally suppressed during oxLDL stress (Groenen, *et al*.,2021). The imbalance between unrestrained lipid influx and diminished capacity of efflux is recognized as the central mechanistic axis that governs foam cell persistence and progression of atherosclerotic lesion. Consequently, therapeutic strategies that are capable for diminishing uptake of oxLDL, curbing oxidative, inflammatory signaling, and cholesterol efflux restoration that are considered highly desirable strategy for anti-atherogenic intervention (Yu *et al*., 2013). Statins, particularly Atorvastatin, remain the cornerstone of clinical management for atherosclerotic cardiovascular disease (Diamantis *et al*., 2017). Beyond their well-established lipid-lowering drug statins exert pleiotropic anti-inflammatory and antioxidant effects on macrophages, including expression of scavenger receptor modulation and expression and cholesterol efflux enhancement contributes to their anti-atherogenic effects (Kouhpeikar *et al*., 2020). Nevertheless, statin therapy entails dose-dependent adverse effects such as myopathy and hepatotoxicity in a certain patient, while residual cardiovascular risk persists despite usage of statin optimally. These limitations have spurred the interest in identifying complementary or alternative agents that are particularly bioactive compounds derived from plants might target foam cell formation cascade with a favorable safety profile.

Medicinal plants are recognized as valuable reservoirs of cardioprotective phytochemicals, and *Aegle marmelos* (Bael), widely used in traditional medicine, has attracted attention for its diverse pharmacological properties, that includes antioxidant, anti-inflammatory, anti-diabetic, and hypolipidemic effects (Ramakrishna *et al*., 2024). Aegeline, an amide alkaloid isolated from the leaves of *Aegle marmelos*, has been reported to exhibit notable bioactivity across several of these domains, and its structural and functional characteristics suggest plausible mechanistic relevance to macrophage lipid handling and redox regulation (Labibah *et al*., 2025). Despite its promise, the specific effects of Aegeline on oxLDL-induced macrophage foam cell formation including scavenger receptor mediated lipid uptake, oxidative stress, inflammatory cytokine release, and cholesterol efflux remains systematically uninvestigated, representing a significant gap in the current understanding of its cardioprotective potential (Lin *et al*., 2017).

Therefore, the present study employed the human THP-1 monocyte/macrophage cell line, a well-validated *in vitro* model for studying foam cell biology, to evaluate the anti-atherogenic potential of Aegeline against oxLDL-induced pathological changes. Particularly, we examined the effects of Aegeline on DiI-oxLDL uptake, pro-inflammatory cytokines release and functional cholesterol efflux capacity (Zheng *et al*., 2017) benchmarking its efficacy against Atorvastatin as a clinically established positive control. By concurrently assessing these interconnected facets of foam cell pathology, this study aims to provide integrated mechanistic evidence supporting the candidacy of Aegeline as a natural lead compound for anti-atherosclerotic therapeutic development and highlighting the potential of Aegeline and atorvastatin in combinatorial therapy for enhanced cardiovascular protection.

## 2.0 Methods

### 2.1 Source of chemicals

Aegeline was procured from Clearsynth Canada. Bovine Serum Albumin (BSA) was procured fromSigma–Aldrich, USA. All other chemicals used were of the analytical grade and were obtained from Medox Biotech India, SRL (Sisco Research Laboratories Pvt. Ltd), Genei, Bangalore and Central Drug House Pvt. Ltd (CDH) Mumbai India). 1,1’-dioctadecyl-3,3,3,3’-tetramethylindocarbocyanine perchlorate oxidized LDL (DiI-oxLDL), Roswell Park Memorial Institute medium (RPMI), Fetal bovine serum (FBS), and Antibiotic-antimycotic (Pen-Strep) were procured from Gibco, Thermo Fisher Scientific Inc. (Waltham, MA, USA). Cholesterol Efflux Assay Kit (ab196985), CD14 Rabbit mab (A19011).

### 2.2 THP-1 cell line procurement and macrophage differentiation

THP1 monocytes were obtained from National Center for Cell Science, Pune, India. The cells are initially grown in RPMI 1640 supplemented with 10% Fetal Bovine Serum (v/v), 100U/ml penicillin-100ug/ml streptomycin in T25 and T75 vented culture flasks. Cultures were incubated at 37◦C in 5% CO_2_/95% humidified air. When the cells reached 80-90% confluence in the flask, they were collected and centrifuged and seeded onto a 96-well, 6-well or 12-well plates. To induce differentiation, THP1 cells were transferred to a differentiation medium (RPMI containing 0.5% FBS, Pen/strep with 100nM Phorbol 12-Myristate 13-Acetate) for 3 days (72 hours) and replenished with fresh 10% complete medium for 2 days (48hours) (Smith *et al*., 2018). Differentiation of THP1 macrophages was validated with Immunofluorescence staining of CD14 THP1 cells. After differentiation, the macrophages were used for further studies.

### 2.3 Validation of THP1 macrophages differentiation-Immunostaining of CD14 marker

To validate the differentiation, THP1 monocytes were cultured on coverslips pre-coated with gelatin 0.2% (in 6 well plates) subjected to 100nM PMA. For staining, the cells were fixed with 4% paraformaldehyde (PFA) for 15minutes and washed thrice with PBS (pH 7.4) and permeabilized with 0.25% Triton X 100 in PBS. After three washes with PBS, the cells were blocked with 5% BSA for 1 h and then incubated with CD14 antibody (1:250, mouse anti-human monoclonal in 5% BSA in PBS) in a light-protected chamber maintained overnight at 4°C. Following incubation, the cells were washed thrice with PBS and incubated with Alexa fluor 594 conjugated secondary antibodies for 1 hour at RT. The cells were then washed three times with PBS and mounted with fluoromount G. Images were acquired with an Accu-scope EXC-500 fluorescence microscope using a 10/40× objective lens

### 2.4 Enzyme linked immunosorbent assay (ELISA)

oxLDL and Inflammatory cytokine levels were assessed using ELISA in the serum samples. Briefly, 100 μL of suitably diluted serum sample containing 25 μg of protein was coated onto 96 well plates using bicarbonate buffer and left overnight at 4°C. The next day, plates were washed with PBS and blocked with 1% BSA solution (200 μl /well) for 2 h to prevent non-specific binding in the subsequent steps. The wells were then washed three times with PBS and dried. The specific primary antibodies suitably diluted in 1% BSA (1:1000) were added to the wells (100μl/well) and incubated overnight at 4°C. The following day, the wells were washed with PBS-Tween-20 four times and incubated with the 100μl of horseradish peroxidase conjugated secondary antibody for an hour at room temperature. After washing thrice with 200μl of PBS, 100 μl of 1mM ABTS (2,2’-Azino-bis (3-ethylbenzothiazoline-6-sulfonic acid)) substrate was added and incubated in the dark for 20 min. Then, 2 N sulfuric acid (100μl/well) was added to the wells to stop the reaction, and the plates were read at 415 nm using a Bio-Rad iMark™ Microplate Absorbance Reader (Bio-Rad Laboratories, Hercules, CA, USA).

### 2.5 Immunofluorescence studies

The differentiated macrophages were treated with DiI-oxLDL for 3 hours at 37◦C and washed with 1ml of PBS thrice. Then treated with Aegeline (), Atorvastatin () and Aegeline, Atrovastatin in combination to DiI-oxLDL exposed cells and incubated at 37◦C for 24hours. After incubation, cells were washed gently with 1ml of PBS thrice. 1μg/μl Hoechst (nuclear stain) was added to counterstain and incubated for 1 minute. The slides were washed again with PBS and mounted with fluoromount G. Images were acquired with an Accu-scope EXC-500 fluorescence microscope using a 10/40× objective lens.

### 2.6 Cholesterol efflux assay

A cholesterol efflux assay kit was used to quantitate the rate of cholesterol efflux using fluorescently labeled cholesterol. Briefly, THP-1 macrophages (1 × 10^5^ cells/mL) were seeded in in 96-well black plates with a clear flat bottom and treated with Aegeline, Atorvastatin and Aegeline + Atorvastatin combination. THP-1 macrophages were exposed to 100 µL of labeling medium mix and incubated for 1 hour in dark at 37 °C in a CO_2_ incubator. The labeling medium was removed and incubated with 100 µL of equilibration medium and incubated overnight (16hours) in dark at 37 °C. After overnight incubation, the medium was aspirated and washed with phenol-red free RPMI medium. After removing the wash medium, 10 µM/mL of Aegeline, 10 µM/mL of Atorvastatin and their combination were incubated for 24hours in dark at 37 °C. At the end of incubation time, supernatant was transferred to a new 96-well black plate, and the remaining cells were lysed by adding 100 μL of the cell lysis buffer and shaking for 30 min at RT. Using a microplate reader (Molecular devices, iD5), fluorescence (Ex/Em = 485/535 nm) was measured. The amount of fluorescence in the supernatant divided by the sum of fluorescence in the cell lysate and supernatant was used to calculate percentage of efflux.

### 2.7 Statistical analysis

All statistical analyses were performed using GraphPad Prism 11.0 software (San Diego, CA, USA). Values are presented as Mean ± Standard error of Mean (SEM). Analysis was done using Two-tailed paired-t-test with Weltch’s corrections and Tukey’s post-hoc test followed by one way ANOVA was performed for multiple group comparison.

## 3.0 Results

### 3.1 Differentiation of THP1 monocytes to macrophages

Initially, to differentiate THP1 monocytes into macrophages, the cells were treated with differentiation medium for 72 hours and left in resting medium for 48 hours **(Figure 1B)**. The macrophage phenotype was confirmed through immunostaining for CD14 a well-established cell surface marker for monocytes **(Figure 1C)**. Compared to the undifferentiated control, the differentiated macrophages showed lower expression of CD14, indicating that the CD14 is expressed to a greater extent in monocytes than the macrophages **(Figure 1C)**.

**Figure 1.**
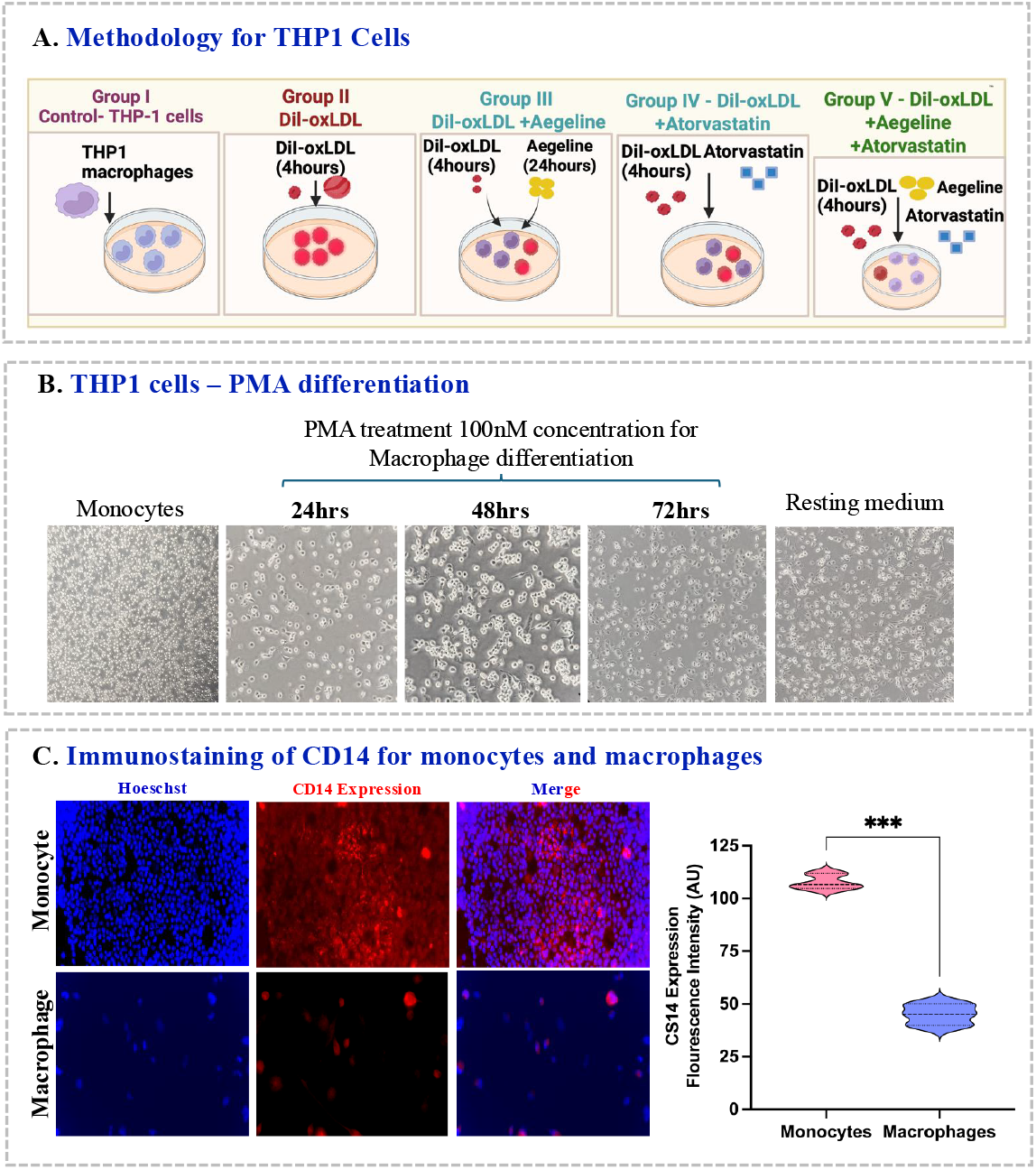
Experimental workflow, THP-1 macrophage differentiation, and phenotypic validation via CD14 immunostaining. **Experimental study design, differentiation of THP-1 monocytes into macrophages, and surface expression of CD14 marker.** **(A)** Schematic representation of the experimental grouping and study workflow, detailing control, DiI-oxLDL and therapeutic interventions (Aegeline, Atorvastatin, and combination treatment). **(B)** Morphological transition during PMA-induced differentiation of human THP-1 monocytes into adherent macrophages. **(C)** Representative immunofluorescence staining depicting CD14 expression (red) and DAPI nuclear counterstain (blue) in differentiated THP-1 macrophages to confirm successful macrophage polarization.

### 3.2 Aegeline and Atorvastatin Suppress oxLDL-Induced Inflammatory Mediators

To evaluate the anti-inflammatory activity of Aegeline, Atorvastatin, and their combination in THP-1 macrophages, key inflammatory markers including C-reactive protein (CRP), tumor necrosis factor-alpha (TNF-α), interleukin-6 (IL-6), and interleukin-8 (IL-8) were quantified using ELISA. Control cells displayed minimal baseline expression across all evaluated inflammatory markers. DiI-oxLDL exposure triggered a robust pro-inflammatory response, marked by prominent upregulation in CRP (p<0.001); TNF-α (p<0.001); IL-6 p<0.001), and IL-8 (p<0.001) secretion when compared to control. Aegeline with oxLDL treated cells attenuated the oxLDL-induced elevation of all four mediators (CRP (p<0.001), TNF-α (p<0.001), IL-6 (p<0.001), IL-8 (p<0.001), when compared to control group. Atorvastatin exerted an overall stronger anti-inflammatory action, significantly reducing TNF-α (p < 0.001), IL-6 (p< 0.01), and IL-8 (p< 0.001). Notably, Atorvastatin potently suppressed CRP expression (p=ns) when compared to control (ns, *p* > 0.05 vs. Control). Combination treatment with Aegeline and Atorvastatin (DiI-oxLDL + Aegeline + Atorvastatin) demonstrated enhanced anti-inflammatory efficacy across multiple cytokines compared to aegeline alone and atoravastatin alone groups.

The combination showed the greatest suppression of TNF-α (p=ns), IL-8 (p<0.01) and markedly attenuated IL-6 (p<0.01) when compared to control and CRP (p<0.001) when compared to control. Collectively, these data demonstrate that both Aegeline and Atorvastatin mitigate oxLDL-induced inflammatory signaling in THP-1 macrophages. While Atorvastatin single-agent treatment is particularly effective at resolving CRP induction, combining Aegeline with Atorvastatin provides superior, broad-spectrum suppression across pro-inflammatory cytokines and chemokines, completely restoring TNF-α expression to physiological levels **(Figure 2A)**.

**Figure 2.**
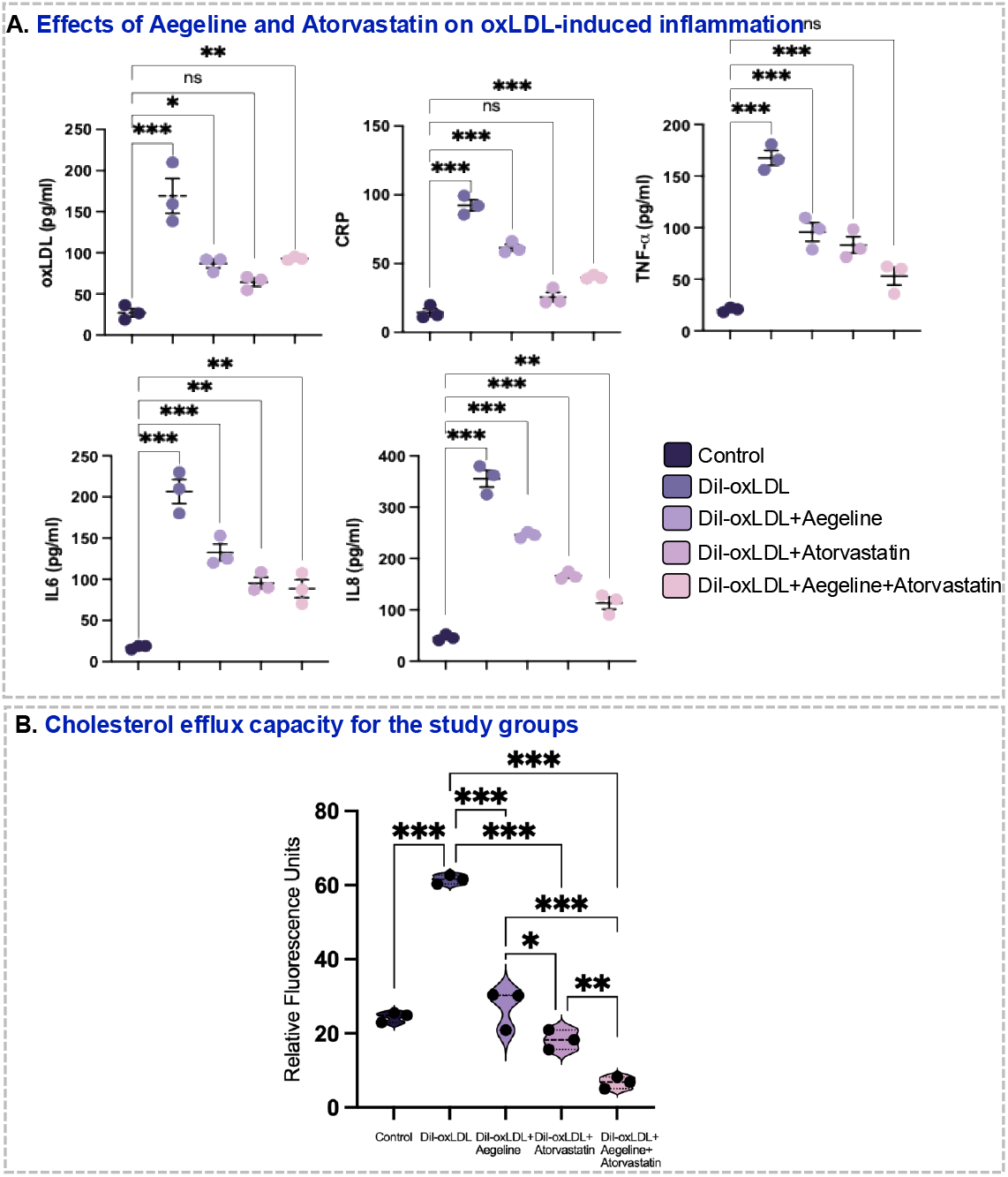
Effect of Aegeline and Atorvastatin treatments on macrophage inflammatory markers and cholesterol efflux. **Effects of Aegeline and Atorvastatin on oxLDL-induced inflammation and cholesterol efflux in macrophages**. (A) Inflammatory panel: Concentrations of oxLDL, CRP, TNF-α, IL6, and IL8 (pg/ml) were quantified under five conditions: Control, Dil-oxLDL, Dil-oxLDL + Aegeline, Dil-oxLDL + Atorvastatin and Dil-oxLDL + Aegeline + Atorvastatin. (B) Cholesterol efflux assay: Relative fluorescence units (RFU) were measured to assess cholesterol efflux across the same five conditions. Violin plots depict distribution, with statistical significance denoted as above. Data are presented as mean ± SEM, with statistical significance indicated by *p < 0.05, **p < 0.01, ***p < 0.001, and ns (non-significant)

### 3.3. Cholesterol efflux capacity for the study groups

Exposure of Aegeline, Atorvastatin and their combination with DiI-oxLDL macrophage cells for 24hours hours significantly reduced the cholesterol uptake compared to untreated cells. Although combinatorial therapy significantly reduces cholesterol uptake relative to the aegeline alone and atorvastatin alone groups, the effect was significantly increased when compared with control cells. Untreated control cells exhibited basal intracellular fluorescence levels. On exposure with DiI-oxLDL significantly increased fluorescence retention (p< 0.001) when compared to control group reflecting impaired efflux and excessive intracellular cholesterol accumulation. On treating with Aegeline (DiI-oxLDL + Aegeline) markedly reduced intracellular cholesterol retention (p < 0.001) when compared to DiI-oxLDL alone group indicating a significant promotion of cholesterol efflux. Atorvastatin treatment (DiI-oxLDL + Atorvastatin) enhanced cholesterol efflux to an even greater extent (p < 0.001) when compared to DiI-oxLDL alone group (p < 0.05). The combination of Aegeline and Atorvastatin (DiI-oxLDL + Aegeline + Atorvastatin) treatment demonstrated the strongest effect, resulting in the lowest residual intracellular cholesterol levels (p< 0.001) when compared to DiI-oxLDL alone groups. These findings indicate that both Aegeline and Atorvastatin reduces the cholesterol uptake in cells, with their combination demonstrating synergistic efficacy in reducing intracellular cholesterol overload **(Figure 2B)**.

### 3.4. Immunofluorescence studies for the effect of Aegeline and Atorvastatin on oxLDL uptake

To evaluate the impact of Aegeline, Atorvastatin, and their combined treatment with DiI-oxLDL, cellular oxLDL accumulation was quantified across experimental groups. Upon DiI-oxLDL exposure, oxLDL levels were increased (p<0.001) when compared to control. While on treatment with aegeline and DiI-oxLDL, noticeably attenuated oxLDL uptake though levels remained modestly elevated compared to the control group (p < 0.05). Treatment with Atorvastatin and DiI-oxLDL demonstrated robust efficacy, reducing oxLDL accumulation to levels that were not statistically significant from the untreated control group (p =ns). The combination treatment of Aegeline and Atorvastatin with DiI-oxLDL similarly reduced oxLDL accumulation relative to DiI-oxLDL alone though values remained significantly higher than baseline control (p< 0.01). Together, these findings indicate that both Aegeline and Atorvastatin effectively suppress oxLDL cellular loading, with Atorvastatin demonstrating the strongest reduction back toward physiological baseline levels **(Figure 3)**.

**Figure 3.**
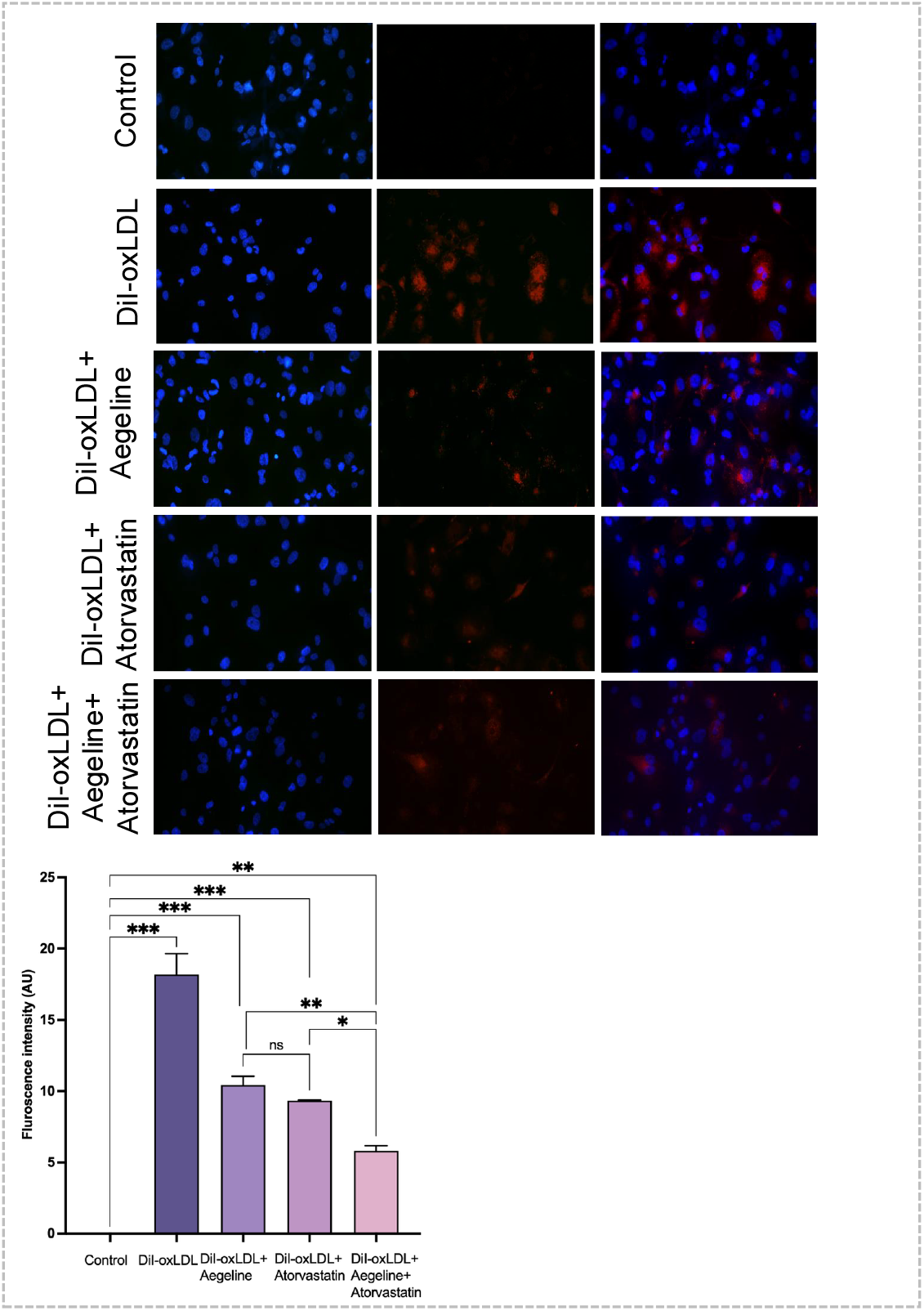
Immunofluorescence quantification of intracellular DiI-oxLDL clearance in THP-1 macrophages. **(A)** Representative Immunofluorescence images at 40x magnification of THP 1 macrophages treated with DiI-oxLDL, Aegeline, Atorvastatin, Aegeline and atorvastatin in combination for assessing the cholesterol efflux capacity of HDL obtained from healthy individuals and patients. **(B)** The fluorescence intensity was measured using using ImageJ software. Values are expressed as mean ± SEM. Statistical analysis was performed using one-way ANOVA with Tukey’s multiple comparisons test \*\**p* < 0.05, \*\**p* < 0.01, \*\**p* < 0.001, between the groups.

## 4.0 Discussion

Atherosclerosis is a chronic inflammatory disorder of the vascular wall characterized by subendothelial retention and oxidation of low-density lipoproteins (oxLDL), impaired cholesterol homeostasis, and persistent macrophage activation (Glass and Witztum, 2001; Tabas and Bornfeldt, 2016). In the early stages of atherogenesis, circulating monocytes infiltrate the arterial intima, differentiate into macrophages, and internalize modified lipoproteins primarily via scavenger receptors (Kunjathoor *et al*., 2002; Mehta and Li, 1998). This unconstrained uptake of oxLDL exceeds the intracellular processing capacity, leading to excessive esterified cholesterol storage within lipid droplets and the generation of foam cells (Goldstein and Brown, 2009; Collins and Cybulsky, 2001). Concurrent with lipid accumulation, oxLDL uptake triggers intracellular oxidative stress and activates redox-sensitive signaling networks, predominantly nuclear factor-kappa B (NF-κB), which drives the transcription of acute-phase proteins, pro-inflammatory cytokines, and chemokines (Collins and Cybulsky, 2001). The resulting local inflammatory milieu impairs intrinsic reverse cholesterol transport mechanisms, exacerbating foam cell entrapment, promoting necrotic core formation, and driving plaque instability (Tall, 2008; Moore and coworkers, 2018). Reversal of lipid overload through the enhancement of macrophage cholesterol efflux represents a crucial therapeutic strategy to attenuate foam cell formation and slow atherosclerotic progression (Cuchel and Rader, 2006). Cholesterol efflux is the rate-limiting initial step of reverse cholesterol transport, wherein intracellular free cholesterol is transported across the plasma membrane to extracellular acceptors (Phillips, 2014; Yvan-Charvet *et al*., 2007). In the present study, challenge with DiI-oxLDL significantly increased intracellular lipid retention and fluorescent oxLDL loading in THP-1 macrophages, confirming severe impairment of baseline efflux capacity. Treatment with Aegeline—a bioactive plant alkaloid isolated from *Aegle marmelos*—substantially enhanced cholesterol efflux and reduced intracellular DiI-oxLDL accumulation. This effect was comparable in magnitude to that observed with Atorvastatin, a 3-hydroxy-3-methylglutaryl-coenzyme A (HMG-CoA) reductase inhibitor well known for upregulating cellular cholesterol export machinery alongside its lipid-lowering actions (Jain and Ridker, 2005)

The mechanisms by which statins promote cholesterol efflux involve both HMG-CoA reductase-dependent pathways and transcriptional upregulation of cholesterol export pathways, mediated in part through nuclear receptors such as peroxisome proliferator-activated receptor gamma (PPAR-γ) and liver X receptor alpha (LXR-α) (Venkateswaran *et al*., 2000). Aegeline has been reported to modulate metabolic pathways and suppress oxidative stress, which may protect cellular export mechanisms from oxidative damage or directly stimulate LXR-α-dependent transactivation (Narender *et al*., 2007). Intriguingly, combining Aegeline with Atorvastatin produced an additive promotion of cholesterol efflux, lowering intracellular DiI-oxLDL retention significantly beyond single-agent treatment. Simultaneous enhancement of cellular export pathways prevents macrophage lipid accumulation and accelerated atherogenesis; thus, dual treatment may synergistically target distinct steps in lipid processing and export trafficking, providing a robust mechanism for clearing accumulated intracellular sterols (Oram and Heinecke, 2005).

In addition to lipid accumulation, oxLDL stimulation in macrophages triggers a potent inflammatory cascade characterized by the upregulation of C-reactive protein (CRP), tumor necrosis factor-alpha (TNF-α), interleukin-6 (IL-6), and interleukin-8 (IL-8) (Libby, 2002).Elevated serum levels of CRP serve as a key clinical biomarker for systemic vascular inflammation and major adverse cardiovascular events (Ridker *et al*., 2002) In our model, oxLDL challenge induced a marked rise in cellular CRP expression, which was partially blunted by Aegeline monotherapy and potently normalized back to baseline levels by Atorvastatin monotherapy. Statins are established to exert direct pleiotropic anti-inflammatory actions independently of lipid lowering, including the down-regulation of acute-phase reactant synthesis through the inhibition of isoprenoid intermediates in the mevalonate pathway (Liao and Laufs, 2005). The pro-inflammatory cytokines TNF-α and IL-6 act in concert to amplify arterial tissue injury, induce endothelial cell adhesion molecules, and sustain systemic immune cell activation. Monotherapy with either Aegeline or Atorvastatin significantly attenuated oxLDL-induced TNF-α and IL-6 secretion, aligning with previous observations that both compounds inhibit upstream NF-κB activation. Notably, combination therapy with Aegeline and Atorvastatin exerted a synergistic inhibitory effect on TNF-α, reducing its concentration back to physiological baseline levels (ns vs. Control).

Similarly, the secretion of IL-8, a critical CXC chemokine responsible for directing neutrophil and leukocyte chemotaxis into the subendothelial space, was markedly elevated following oxLDL loading. Both Aegeline and Atorvastatin reduced IL-8 expression, with the combined treatment demonstrating superior suppression compared to monotherapies. Elevated intracellular cholesterol directly promotes Toll-like receptor (TLR) assembly and signaling in membrane lipid rafts; conversely, enhancing cholesterol efflux disrupts lipid raft architecture and suppresses downstream chemokine secretion. Therefore, the enhanced clearance of intracellular oxLDL mediated by dual Aegeline and Atorvastatin treatment directly reinforces their combined anti-inflammatory efficacy, establishing a favorable feedback loop between lipid clearance and inflammation resolution.

## 5.0. Conclusion

In conclusion, these findings demonstrate that Aegeline functions as a potent bioactive agent capable of improving cholesterol efflux and suppressing oxLDL-induced inflammatory responses in THP-1 macrophages. While Atorvastatin exhibits high single-agent efficacy in restoring CRP levels, combining Aegeline with Atorvastatin offers complementary, multi-target therapeutic benefits that are most notably by maximizing reverse cholesterol transport and completely resolving TNF-α production back to baseline levels. Further studies in animal models of atherosclerosis are warranted to confirm whether this dual therapeutic approach effectively stabilizes vulnerable plaques and halts atheroma progression.

## Author contributions

**<u>Abinayaa Rajkumar:</u>** Conceptualization (First); data curation (First); formal analysis (First); investigation (First); methodology (First); visualization (First); writing − original draft (First); writing − review and editing (First). **<u>Cathline Mary Ramesh:</u>** Conceptualization; data curation; formal analysis; Visualization and writing − review and editing (supporting). **<u>Ezhila Subramani DhatchanamoorthyVedhanayaki</u>**: Visualization and writing − review and editing (supporting). **<u>Kalaiselvi Periandavan</u>:** Conceptualization (lead); funding acquisition (lead); project administration (lead); resources (lead); supervision (lead); validation (lead); visualization (lead); writing − review and editing (lead). All authors have read and approved the final version of the manuscript.

## Acknowledgements

The authors express their gratitude to the financial support provided to Cathline Mary Ramesh by **Rashtriya Uchchatar Shiksha Abhiyan (RUSA 2.0)**, Research, Innovation and Quality Improvement, University of Madras in the form of Project Fellow. The authors also acknowledge **Department of Health Research - Multidisciplinary Research Unit (DHR-MRU)** at the Dr. ALM Post Graduate Institute for Basic Medical Sciences, University of Madras, for providing access to its state-of-the-art facilities.

## Data Availability Statement

The data supporting the findings of this study are available from the corresponding author upon reasonable request.

## Declaration of Transparency and Scientific Rigor

This Declaration acknowledges that this paper adheres to the principles for transparent reporting and scientific rigor of preclinical research recommended by funding agencies, publishers, and other organizations engaged in supporting research.

## References

Björkegren, J. L., & Lusis, A. J. (2022). Atherosclerosis: recent developments. Cell, 185(10), 1630–1645.

Borén, J., Packard, C. J., & Binder, C. J. (2025). Apolipoprotein B-containing lipoproteins in atherogenesis. Nature Reviews Cardiology, 22(6), 399–413.

Brand, K., Page, S., Rogler, G., Bartsch, A., Brandl, R., Knuechel, R., … & Neumeier, D. (1996). Activated transcription factor nuclear factor-kappa B is present in the atherosclerotic lesion. The Journal of clinical investigation, 97(7), 1715–1722.

Brand, K., Page, S., Walli, A. K., Neumeier, D., & Baeuerle, P. A. (1997). Role of nuclear factorLJkappa B in atherogenesis. Experimental Physiology: Translation and Integration, 82(2), 297–304.

Collins, T., & Cybulsky, M. I. (2001). NF-kappaB: pivotal mediator or innocent bystander in atherogenesis?. The Journal of clinical investigation, 107(3), 255–264. 10.1172/JCI10373

Cuchel, M., & Rader, D. J. (2006). Macrophage reverse cholesterol transport: key to the regression of atherosclerosis?. Circulation, 113(21), 2548–2555. 10.1161/CIRCULATIONAHA.104.475715

Diamantis, E., Kyriakos, G., Victoria Quiles-Sanchez, L., Farmaki, P., & Troupis, T. (2017). The anti-inflammatory effects of statins on coronary artery disease: an updated review of the literature. Current cardiology reviews, 13(3), 209–216.

Falk, E. (2006). Pathogenesis of atherosclerosis. Journal of the American College of cardiology, 47(8S), C7–C12.

Glass, C. K., & Witztum, J. L. (2001). Atherosclerosis. the road ahead. Cell, 104(4), 503–516. 10.1016/s0092-8674(01)00238-0

Goldstein, J. L., & Brown, M. S. (2009). The LDL receptor. Arteriosclerosis, thrombosis, and vascular biology, 29(4), 431–438. 10.1161/ATVBAHA.108.179564

Groenen, A. G., Halmos, B., Tall, A. R., & Westerterp, M. (2021). Cholesterol efflux pathways, inflammation, and atherosclerosis. Critical reviews in biochemistry and molecular biology, 56(4), 426–439.

Jain, M. K., & Ridker, P. M. (2005). Anti-inflammatory effects of statins: clinical evidence and basic mechanisms. Nature reviews. Drug discovery, 4(12), 977–987. 10.1038/nrd1901

Jebari-Benslaiman, S., Galicia-García, U., Larrea-Sebal, A., Olaetxea, J. R., Alloza, I., Vandenbroeck, K., … & Martín, C. (2022). Pathophysiology of atherosclerosis. International journal of molecular sciences, 23(6), 3346.

Kleinbongard, P., Heusch, G., & Schulz, R. (2010). TNFα in atherosclerosis, myocardial ischemia/reperfusion and heart failure. Pharmacology & therapeutics, 127(3), 295–314.

Kouhpeikar, H., Delbari, Z., Sathyapalan, T., Simental-Mendía, L. E., Jamialahmadi, T., & Sahebkar, A. (2020). The effect of statins through mast cells in the pathophysiology of atherosclerosis: a review. Current Atherosclerosis Reports, 22(5), 19.

Kunjathoor, V. V., Febbraio, M., Podrez, E. A., Moore, K. J., Andersson, L., Koehn, S., Rhee, J. S., Silverstein, R., Hoff, H. F., & Freeman, M. W. (2002). Scavenger receptors class A-I/II and CD36 are the principal receptors responsible for the uptake of modified low density lipoprotein leading to lipid loading in macrophages. The Journal of biological chemistry, 277(51), 49982–49988. 10.1074/jbc.M209649200

Labibah, R., Bilqis, A. A., Tanzil, P. J. O., Diarsvitri, W., Rahadianto, R., & Wisanti, R. (2025). The effect of the combination of ethanolic extracts of bael leaves (Aegle marmelos) and brown algae (Padina australis) on HDL levels in Rattus norvegicus fed a high-fat diet. MEDISAINS: Jurnal Ilmiah Ilmu-Ilmu Kesehatan, 23(2), 106–110.

Leiviskä, J., Sundvall, J., Alfthan, G., Tähtelä, R., Salomaa, V., Jauhiainen, M., & Vartiainen, E. (2013). What have we learnt about high-density lipoprotein cholesterol measurements during 32 years? Experiences in Finland 1980–2012. Clinica Chimica Acta, 415, 118–123.

Li, Y., Shen, S., Ding, S., & Wang, L. (2018). Toll-like receptor 2 downregulates the cholesterol efflux by activating the nuclear factor-κB pathway in macrophages and may be a potential therapeutic target for the prevention of atherosclerosis Retraction in/10.3892/etm.2024.12760. Experimental and Therapeutic Medicine, 15(1), 198–204.

Liao, J. K., & Laufs, U. (2005). Pleiotropic effects of statins. Annual review of pharmacology and toxicology, 45, 89–118. 10.1146/annurev.pharmtox.45.120403.095748

Libby P. (2002). Inflammation in atherosclerosis. Nature, 420(6917), 868–874. 10.1038/nature01323

Lin, X. L., Hu, H. J., Liu, Y. B., Hu, X. M., Fan, X. J., Zou, W. W., … & Gu, C. H. (2017). Allicin induces the upregulation of ABCA1 expression via PPARγ/LXRα signaling in THP-1 macrophage-derived foam cells. International journal of molecular medicine, 39(6), 1452–1460.

Maxfield, F. R., & Tabas, I. (2005). Role of cholesterol and lipid organization in disease. Nature, 438(7068), 612–621. 10.1038/nature04399

Mehta, J. L., & Li, D. Y. (1998). Identification and autoregulation of receptor for OX-LDL in cultured human coronary artery endothelial cells. Biochemical and biophysical research communications, 248(3), 511–514. 10.1006/bbrc.1998.9004

Moore, K. J., & Tabas, I. (2011). Macrophages in the pathogenesis of atherosclerosis. Cell, 145(3), 341–355.

Moore, K. J., Sheedy, F. J., & Fisher, E. A. (2013). Macrophages in atherosclerosis: a dynamic balance. Nature reviews. Immunology, 13(10), 709–721. 10.1038/nri3520

Narender, T., Shweta, S., Tiwari, P., Papi Reddy, K., Khaliq, T., Prathipati, P., Puri, A., Srivastava, A. K., Chander, R., Agarwal, S. C., & Raj, K. (2007). Antihyperglycemic and antidyslipidemic agent from Aegle marmelos. Bioorganic & medicinal chemistry letters, 17(6), 1808–1811. 10.1016/j.bmcl.2006.12.037

Oram, J. F., & Heinecke, J. W. (2005). ATP-binding cassette transporter A1: a cell cholesterol exporter that protects against cardiovascular disease. Physiological reviews, 85(4), 1343–1372. 10.1152/physrev.00005.2005

Pamukcu, B., Lip, G. Y., & Shantsila, E. (2011). The nuclear factor–kappa B pathway in atherosclerosis: a potential therapeutic target for atherothrombotic vascular disease. Thrombosis research, 128(2), 117–123.

Phillips M. C. (2014). Molecular mechanisms of cellular cholesterol efflux. The Journal of biological chemistry, 289(35), 24020–24029. 10.1074/jbc.R114.583658

Ramakrishna, S., & Gn, V. (2024). A comprehensive review on traditional uses, phytochemistry, and pharmacological activities of Aegle marmelos. Journal of Pharma Insights and Research, 2(2), 136–142.

Ridker, P. M., Rifai, N., Rose, L., Buring, J. E., & Cook, N. R. (2002). Comparison of C-reactive protein and low-density lipoprotein cholesterol levels in the prediction of first cardiovascular events. The New England journal of medicine, 347(20), 1557–1565. 10.1056/NEJMoa021993

Steinberg D. Atherosclerosis: basic mechanisms. Oxidation, inflammation, and genetics. Circulation. 1995;91(9):2488–2496. doi:10.1161/01.CIR.91.9.2488.

Tabas, I., & Bornfeldt, K. E. (2016). Macrophage Phenotype and Function in Different Stages of Atherosclerosis. Circulation research, 118(4), 653–667. 10.1161/CIRCRESAHA.115.306256

Tall, A. R. (2008). Cholesterol efflux pathways and other potential mechanisms involved in the atheroLJprotective effect of high density lipoproteins. Journal of internal medicine, 263(3), 256–273.

Venkateswaran, A., Laffitte, B. A., Joseph, S. B., Mak, P. A., Wilpitz, D. C., Edwards, P. A., & Tontonoz, P. (2000). Control of cellular cholesterol efflux by the nuclear oxysterol receptor LXR alpha. Proceedings of the National Academy of Sciences of the United States of America, 97(22), 12097–12102. 10.1073/pnas.200367697

Yu, X. H., Fu, Y. C., Zhang, D. W., Yin, K., & Tang, C. K. (2013). Foam cells in atherosclerosis. Clinica chimica acta, 424, 245–252.

Yvan-Charvet, L., Ranalletta, M., Wang, N., Han, S., Terasaka, N., Li, R., Welch, C., & Tall, A. R. (2007). Combined deficiency of ABCA1 and ABCG1 promotes foam cell accumulation and accelerates atherosclerosis in mice. The Journal of clinical investigation, 117(12), 3900–3908. 10.1172/JCI33372

Zheng, S., Du, Y., Ye, Q., Zha, K., & Feng, J. (2021). Atorvastatin enhances foam cell lipophagy and promotes cholesterol efflux through the AMP-activated protein kinase/mammalian target of rapamycin pathway. Journal of cardiovascular pharmacology, 77(4), 508–518.

